# Binary Discriminator Facilitates GPT-based Protein Design

**DOI:** 10.1101/2023.11.20.567789

**Authors:** Zishuo Zeng, Rufang Xu, Jin Guo, Xiaozhou Luo

## Abstract

Generative pre-trained transformers (GPT) models provide powerful tools for de novo protein design (DNPD). GPT-based DNPD involves three procedures: a) finetuning the model with proteins of interest; b) generating sequence candidates with the finetuned model; and c) prioritizing the sequence candidates. Existing prioritization strategies heavily rely on sequence identity, undermining the diversity. Here, we coupled a protein GPT model with a custom discriminator, which enables selecting candidates of low identity to natural sequences while highly likely with desired functions. We applied this framework to creating novel antimicrobial peptides (AMPs) and malate dehydrogenases (MDHs). Experimental verification pinpointed four broad-spectrum AMPs from 24 candidates. Comprehensive computational analyses on the prioritized MDHs candidates provide compelling evidence for the anticipated function. During experimental validation, 4/10 and 3/10 natural MDHs and generated-prioritized novel candidates, respectively, were expressed and soluble. All the soluble candidates (3/3) are functional in vitro. This framework is time- and data-efficient and may therefore considerably expedite the DNPD process.

## Introduction

*de novo* protein design (DNPD), i.e., generating artificial proteins with desired functions, has proven to be a powerful tool in overcoming bottlenecks across various fields, including drug discovery ^1–3^, biosensor engineering ^4–6^, and synthetic biology ^7–9^. DNPD is being accelerated by recent advancements in generative large language models (LLMs, including GPT) ^10^, for examples, ProtGPT2 ^11^ and ProGen ^12^. Similar to the LLMs for natural language processing (NLP) ^13^, these protein generative models were trained on large amount of “text corpus” (proteomes) and can generate “sentences” (protein sequences) that captures the “language semantics” (physicochemical and thermodynamic properties similar to natural proteins). Besides, just like LLMs for NLP can be finetuned to adapt to a specific task (such as automatic dialogues ^14^, summarizing text ^15^, and poetry writing ^16^), protein generative models can be finetuned ^17^ to generate artificial proteins bearing a specific function of interest. Therefore, protein LLM holds great promise in DNPD.

However, evaluating the quality of the generated sequences remains a challenge, as protein sequences are not human-understandable like natural languages. In the work of ProGen ^12^, evaluating the generated sequences was mainly done by per-token log-likelihood scoring, which computes the per-residue conditional probabilities along the sequence, conceptually akin to perplexity, which measures the model’s confidence in the generated sequence. However, limitations of perplexity in the NLP field have been noticed, such as sensitivity to text length and repeated content ^18^, inability to reflect semantic coherence ^19^, and disagreements with human judgements ^20^. Another common approach for sequence evaluation is through its identity to known natural proteins of interest, which was used in a generative adversarial network (GAN)-based framework, ProteinGAN ^21^. In this approach, the generated sequences are prioritized based on their similarity to the known homologs. Both per-token log-likelihood and sequence identity may overlook the promising (likely to possess the desired protein function) but novel (very different from known natural proteins) sequences – the dark matters in the potential proteome universe.

In addition to per-token log-likelihood and sequence identity, a variety of existing protein predictors may be used to prioritizing the generated protein candidates from different perspectives, such as protein structure ^22^ and enzyme commission (EC) number ^23^. Johnson et al. also incorporated multiple sequence-based and structure-based tools (e.g., Rosetta ^24^, ESM-v1 ^25^, CARP-640M ^26^, ProteinMPNN ^22^, MIF-ST ^27^, and ESM-IF ^28^) for predicting whether the generated enzymes possess *in vitro* activity ^29^. However, using these existing predictors suffer from multiple limitations. First, none of these predictors directly reports the likelihood of the desired functions. Second, it requires non-trivial efforts in benchmarking and programming environment setup. Third, these predictors may not be applicable to different types of proteins (e.g., EC number prediction only applies to enzymes).

To address the limitations of current prioritization strategies, here we coupled ProtGPT2 with a convolutional neural network (CNN)-based protein discriminator. Positive protein data are used to finetune the protein generator, ProtGPT2; while both positive and negative data are used to train the discriminator, which will be used to prioritize the generated sequence candidates, allowing a one-stop solution to DNPD for different types of proteins without the dependencies on external tools. A pre-trained LLM, ProtTrans, was used for protein feature extraction for the discriminator. Such LLM-based contextualized embedding technique has been proven to outperform traditional protein features such as one-hot encoding ^30–32^. The efficacy of the framework was then demonstrated by two distinct types of sequences: AMPs as representatives of short sequences lacking homology, and malate dehydrogenases (MDHs) as representatives of longer sequences with inferable homology. In addition, the prompts length and size of finetuning set have been investigated to fully elucidate the potential of this framework. Altogether, our results position this framework as a powerful and efficient tool to advance DNPD.

## Materials and Methods

### Data preparation

We downloaded 3,011 antibacterial peptides from SATPdb database ^33^ (http://crdd.osdd.net/raghava/satpdb/, accessed on 8/24/2022) as positive set. We filtered this peptide set to only include those composed of canonical amino acids and that were less than or equal to 50 residues in length, resulting in 2,879 peptides, which we refer to as the AMP set. For the negative set, we downloaded 13,225 short (less than or equal to 50-residue-long) protein sequences/peptides from UniProt ^34^ (https://www.uniprot.org/, accessed on 8/16/2022). We removed sequences containing certain keywords (“antimic*”, “antibac*”, “*toxic*”, “defensive”) from the UniProt description. This negative set curation is consistent with other relevant methods ^35^ To create a balanced negative set and to avoid bias in sequence length distribution, we randomly sampled from this UniProt pool strictly following the AMP set’s length distribution. For example, if there were 179 AMPs that are 21-residue-long, then we sampled exactly 179 sequences that are 21-residue-long. In total, we collected 2,879 non-AMPs as negative set.

We gathered a positive set of 16,706 MDH sequences from the training set of Repecka et al.’s study ^21^. For negative set, we searched “enzyme” on UniProt (https://www.uniprot.org/) and downloaded all reviewed sequences (n=568,002, accessed on 8/16/2022). To later evaluate how well our model discriminates MDH from functionally close enzymes, we used the validation set (n=213) from Repecka et al. as positive set; we also identified 5,766 EC1.1.1.X sequences (first three enzyme commission [EC] ^36^ levels are 1.1.1 and the last level is not 37) and 543 EC1.1.X.X sequences (first three EC levels are 1.1 and the third level is not 1) as negative set. We then narrowed these sequences to those a) whose EC numbers are not 1.1.1.37 (the EC number of MDH); b) whose sequence lengths are within the range of the positive set’s (64-505); and c) that are not functionally close (EC1.1.1.X or EC1.1.X.X), resulting in 206,596 non-MDH sequences as negative set.

To compile a balanced negative set without bias in sequence length distribution, we enumerated the number of sequences by an interval of 50 and sampled that amount of non-MDH sequences for that interval. For example, if there were 39 MDH sequences that are 110-160 residues long, then we sampled 39 non-MDH sequences of that length range. As a result, we collected 16,706 non-MDH sequences as negative set.

Since the 16,706 MDH sequences may vary a lot (with a length range of [64-505]), to ensure consistency during the finetuning process, we clustered the MDH sequences into multiple clusters using cd-hit ^37^. To reduce the number of resulting clusters, we used the lowest allowed sequence identity threshold (0.4) and word length (2) for clustering. A total of 82 clusters were generated and the majority of these clusters are singletons or very small. The four largest clusters constituted 97.3% (n=16,258) of all MDH sequences. We termed these four clusters as *cluster765*, *cluster2029*, *cluster5987*, and *cluster7477*, with the number inside the name indicating the number of sequences in that cluster (e.g., *cluster765* contain 765 sequences).

### Generator finetuning and sample generation

For the AMP generation task, we finetuned ProtGPT2 ^11^ (https://huggingface.co/nferruz/ProtGPT2) with default parameters on the AMP set. We then generated 3,000 sequences using text generation pipelines from Python (v. 3.8.15) transformers package (v. 4.26.0.dev0) ^38^ (https://huggingface.co/docs/transformers/index) with maximum lengths being 20. Note that the generated sequences may exceed this maximum length. As negative control for the finetuning process, we also generated 3,000 sequences using the non-finetuned ProtGPT2 model.

For the MDH generation task, we finetuned the ProtGPT2 model using four separate datasets: *cluster765*, *cluster2029*, *cluster5987*, and *cluster7477*, resulting in four finetuned models. To generate new sequences, we supplied each finetuned model with a starter sequence of varying length, ranging from 0-100 residues, taken from the representative sequence of the corresponding finetuning set. The maximal length for generation is set to be 400. The results of this sample generation are summarized in Table 1 below. *All finetuning tasks were conducted on* NVIDIA A100 (80G) graphical processing unit (GPU).

**Table 1.** Summary of the MDH sequence generation.

| Group name | Group meaning | Template | Starter seq. length | No. generated seq. |
| --- | --- | --- | --- | --- |
| random | randomly generated according to | representative seq. of | 0 | 1000 |
|  | the amino acid occurrence<br>distribution of all MDH | *cluster7477* | 10 | 1000 |
|  |  |  | 50 | 1000 |
|  |  |  | 100 | 1000 |
| noft | generated by non-finetuned<br>ProtGPT2 | representative seq. of<br>*cluster7477* | 0 | 1000 |
|  |  |  | 10 | 1000 |
|  |  |  | 50 | 1000 |
|  |  |  | 100 | 1000 |
| *cluster765* | finetuned by *cluster765* | representative seq. of<br>*cluster765* | 0 | 1000 |
|  |  |  | 10 | 1000 |
|  |  |  | 50 | 1000 |
|  |  |  | 100 | 1000 |
| *cluster2029* | finetuned by *cluster2029* | representative seq. of<br>*cluster2029* | 0 | 1000 |
|  |  |  | 10 | 1000 |
|  |  |  | 50 | 1000 |
|  |  |  | 100 | 1000 |
| *cluster5987* | finetuned by *cluster5987* | representative seq. of<br>*cluster5987* | 0 | 1000 |
|  |  |  | 10 | 1000 |
|  |  |  | 50 | 1000 |
|  |  |  | 100 | 1000 |
| *cluster7477* | finetuned by *cluster7477* | representative seq. of<br>*cluster7477* | 0 | 1000 |
|  |  |  | 10 | 1000 |
|  |  |  | 50 | 1000 |
|  |  |  | 100 | 1000 |

### Discriminator training

For the AMP discriminator, we split the balanced AMP/non-AMP set (n=2,879+2,879) into training, validation, and testing set in 8:1:1 ratio. From the MDH/non-MDH balanced set (n=16,706+16,706), we randomly sampled 4,500 sequences for training (13.47%) and 500 sequences for validation (1.50%), and the remaining 28,412 sequences were used for testing (85.03%). We performed discriminator training, hyperparameter tuning, and performance evaluation on training set, validation set, and testing set, respectively.

To extract features from the protein sequences, we utilized the ProtT5-XL-UniRef50 model, which was pretrained and publicly available in the ProtTrans project ^39^. We chose ProtT5-XL-UniRef50 among others in ProtTrans because it achieved the best performance most often in different predictive tasks ^39^. We used the transformers package (v. 4.26.0.dev0) in Python (v. 3.8.15) to implement the embedding, with maximum lengths of 50 for the AMP task and 505 for the MDH task.

For the discriminator architecture, we used Keras package (v. 2.10.0) ^40^ to implement a convolutional neural network (CNN) followed by a fully connected neural network (FCNN) (Figure 1). Each convolution layer was coupled with a maximum-pooling layer, and the last maximum-pooling layer was flattened, which was fed to the FCNN. Recticfied linear unit ^41^ was used as activation function for all layers except for the output layer, where the activation function was sigmoid function^42^ instead. We then used Adam ^43^ as the optimizer, binary cross-entropy (Eqn. 1) as the loss function, and accuracy (Eqn. 2) as the evaluation metric. We set maximum number of training epochs to be 100 with early stopping (patience=5). We adopted the following hyperparameter setup after tuning: three 3×3 convolution filter, each followed by 2×2 max-pooling layer; a flatten layer; three fully connected layers (with 1024, 512, 256 nodes, respectively) and one output layer. Tunable hyperparameters include number of filters in CNN, kernel size in CNN, maximum-pooling size, number of dense layers in FCNN, number of nodes in each FCNN layer, batch size, and learning rate of optimizer. Embedding and training were carried out on a NVIDIA A100 (80G) graphical processing unit.

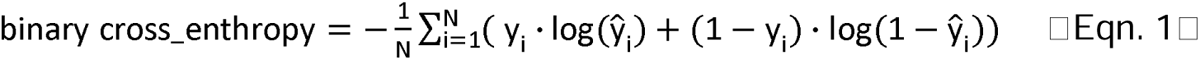

**Figure 1.**
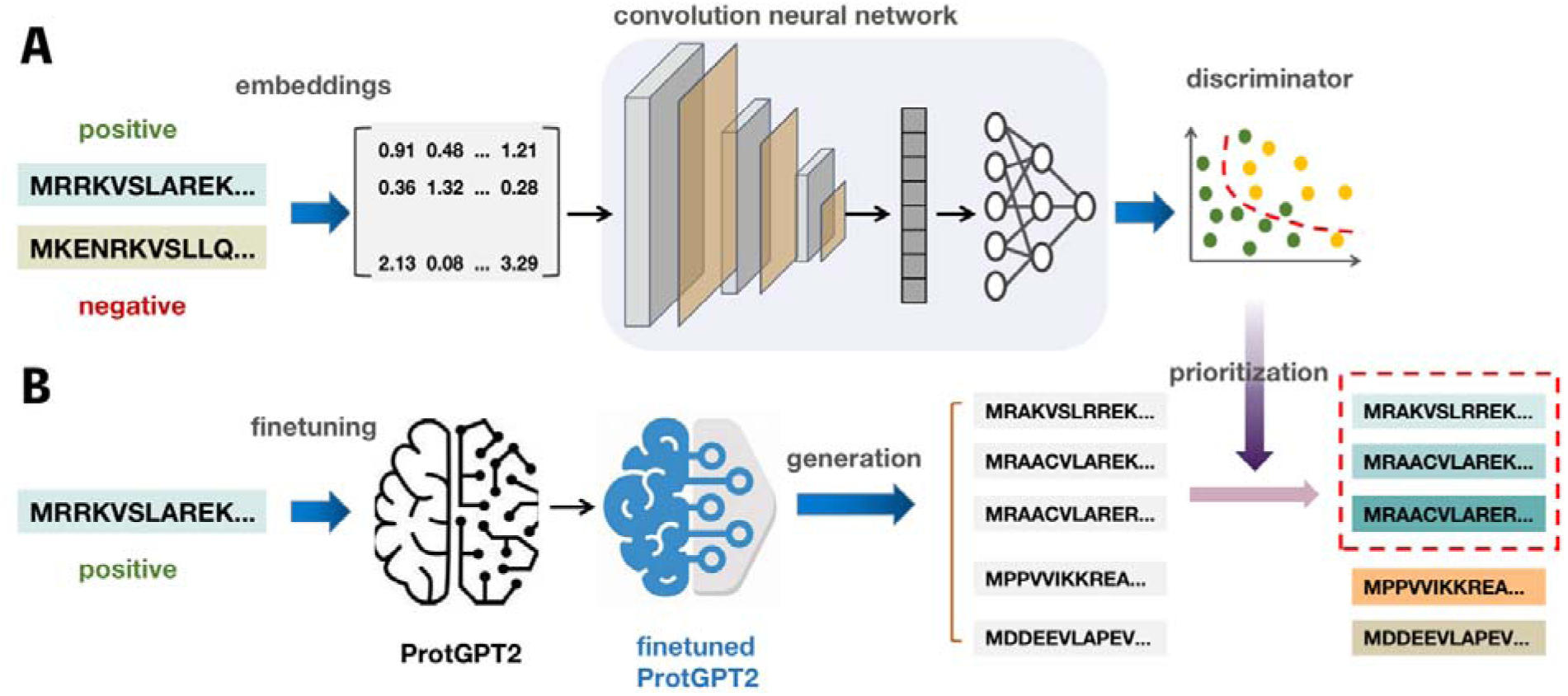
Discriminator (A) and generator (B) architecture. In panel A, positive and negative sequences are embedded using ProtTrans (each residue is converted to a 1,024-value long vector). The embedded inputs are fed to a neural network with three convolution layers and three max-pooling layers, followed by a fully connected neural network. Meanwhile, in panel B, the positive sequences are used to finetune ProtGPT2, which is then used to generate positive candidate sequences. In the end, the discriminator is used to screen the most likely positive sequence candidates.

.where N is the number of samples, y_1_ and 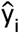 are the *i-*th actual and predicted label, respectively.

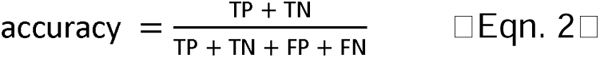

where *TP*, *TN*, *FP*, and *FN* are true positive, true negative, false positive, and false negative, respectively.

### Bioinformatics analysis

We performed Pfam domain ^44^ scanning on all generated MDH sequences using HMMER package^45^ (v. 3.3.2). Pfam profiles were built using hmmerbuild command with Pfam seed alignment flatfile and and Pfam profile flatfile (downloaded from http://ftp.ebi.ac.uk/pub/databases/Pfam/current_release/Pfam-A.seed.gz and http://ftp.ebi.ac.uk/pub/databases/Pfam/current_release/Pfam-A.hmm.gz, respectively. accessed on 11/10/2022). The E-value threshold for Pfam domain scanning was set as 0.001.and we considered a sequence to have MDH’s signature domain if at least one “lactate/malate dehydrogenase” or “Malate/L-lactate dehydrogenase” was identified.

We evaluated the sequence redundancy within *cluster765*, *cluster2029*, *cluster5987*, *cluster7477*, as well as the generated MDH sequences using cd-hit (parameters set as default). We defined “ratio of non-redundant sequences” as the number of non-redundant sequences determined by cd-hit divided by the number of input sequences.

For every generated MDH sequence, we performed pairwise alignment using BLAST standalone package ^46^ (v. 2.13.0+) against all known MDH sequences (n=16,706) and identified the “best hit” (highest bit score) sequence. We recorded the identity between query (generated) sequences and their corresponding best hit sequence. Based on these best hit sequences’ identity and ungapped alignment length, we utilized HFSP curve (homology-derived functional similarity of proteins ^47^, HFSP=0) to visualize whether a generated sequence share the same function as its best hit known MDH.

### Sequence space visualization using t-SNE

We randomly selected 100 known MDHs from *cluster2029*, 100 MDH candidates generated by *cluster2029*-finetuned ProtGPT2 that are prioritized (non-redudant and discriminator prediction>0.95), 100 EC1.1.1.X and 100 EC1.1.X.X for visualization of sequence spatial distribution with t-distributed stochastic neighbor embedding (t-SNE) algorithm ^48^. Each of the selected sequences were embedded by the ProtT5-XL-UniRef50 model ^39^ as an *L*×1024 matrix (where *L* is the length of the protein), which was then flattened into a *L*×1024-long vector as protein-level features. The t-SNE transformation from the embedded features to two-dimensional space was conducted using Python scikit-learn package (v1.1.2) ^49^.

### Mapping conservation to enzyme functionally important sites

Finetuned-generated MDH candidates that are a) non-redundant and b) with discriminator prediction>0.95 were selected. We used a known MDH sequence (UniProt ID: P11708) with determined protein structure (protein data bank [PDB] ^50^ ID: 4MDH) as the reference sequence and was BLASTed against the selected MDH candidates. From the selected MDH candidates with significant hit, 499 were randomly sampled and then subjected to multiple sequence alignment (MSA) along with P11708 using MAFFT ^51^ program. The MSA-derived entropy scores were computed using the msa package ^52^ in R. The conservation scores were calculated as the maximal entropy minus the actual entropy at a given position, so that higher score indicates higher conservation. Annotation of functionally important sites (substrate binding, NAD binding, and sulfate binding) of 4MDH were obtained from PDB. Conservation scores were mapped to residue positions in 4MDH. Visualization of 4MDH was performed using PyMOL ^53^.

### Statistical analysis and visualization

Statistical analysis and visualization were conducted in R (v. 4.2.2) with ggplot2 package (v. 3.4.0). Calculation for area under the receiver operating characteristic curve (ROC-AUC) was done with pROC package ^54^ (v. 1.18.0).

### Experimental validation of AMP candidates

Solid-phase peptide synthesis was used to synthesize crude peptides (which contained 40-70% of the target peptide) by GenScript Biotech in Nanjing, China. Mass spectrometry was used to determine their precise molecular weights. The peptides used for the minimum inhibitory concentration (MIC) were further purified to a 90% degree of purity, as determined by high-performance liquid chromatography.

We used six strains of bacteria in our experiment: four Gram-positive strains (*Staphylococcus aureus* GDMCC 1.221, *Bacillus pumilus* GDMCC 1.225, *Bacillus subtilis* GDMCC 1.222, and *Kocuria rhizophila* GMDCC 1.226) and two Gram-negative strains (*Escherichia coli* GDMCC 1.335 and *E. coli* DH5α). All strains were obtained from Guangdong Microbial Culture Collection Center in Guangzhou, China, except for *E. coli* DH5α, which was maintained in our lab. Each bacterial strain was streaked on Luriae-Bertani (LB) agar medium and incubated overnight at the optimum growth temperature according to the provider’s instructions. Individual colonies were picked and inoculated into LB culture medium, which was then shaken at 200 rpm at the optimum growth temperature overnight. The bacterial suspension was diluted 1:100 using fresh LB culture medium and then cultured for 4-5 hours to reach the logarithmic phase (OD600 of 0.4-0.6). The suspension was adjusted to OD600=0.1 by LB solution and then diluted 1,000 times. A volume of 100 μL cell suspension was transferred into 96-well plates for the next test.

For the bacterial inhibition experiment, we used crude peptides with a target peptide content of approximately 50% for ease of calculation. Freeze-dried peptide powders were dissolved in dimethylsulfoxide (DMSO) to a concentration of 3 mM target peptide. For the antibacterial activity test, peptides (or an equal volume of DMSO solution as a negative control) were added to 96-well plates (each well containing 100 μL cell suspension). The peptide solution and fresh LB solution were adjusted to reach a concentration of 60 μM. Blank controls were set up using LB solution without peptide/DMSO. OD600 values were measured after shaking at 500 rpm at the optimum growth temperature for 12 hours using a high-speed oscillating incubator (ZQZY-88BHS, Shanghai Zhichu Instrument Co., Ltd., China). The blank control OD600 value was used for data normalization. Dunnett’s test was used to compare experimental groups with the control group (two-sided) using OriginPro software (version 9.2, 64 bit). All experiments were performed with three independent replicates.

The peptides used for the MIC determination were more than 90% pure. We prepared a series of peptide concentrations ranging from 0.94 μM to 120 μM. The same culture steps were conducted as described above, except that the final incubation step lasted for 16-18 hours. MICs were determined as the minimum concentration of AMPs where bacteria showed no detectable growth. All experiments were performed with three independent replicates.

### Experimental validation of the MDH candidates

To evaluate the performance of our framework in generating functional enzymes, we randomly selected 10 MDH candidates from a pool of candidates that are of high discriminator prediction (>0.9) and low identity (<50%) to any natural MDHs included in ProteinGAN ^21^. As positive controls, we also randomly selected 10 natural MDHs from the training set of ProteinGAN ^21^.

Sequences were synthesized by Tsingke with codon optimization for expression in *E. coli* BL21 (DE3), and subcloned into a pET-21a(+) plasmid with a His-tag at the 3’ end. The integrity of the obtained constructs was confirmed via DNA sequencing. Plasmids encoding the investigated MDHs were transformed into chemically competent *E. coli* BL21 (DE3) cells. A single colony from a transformant agar plate was picked and incubated in Luria Broth (LB) medium with 100 μg/mL ampicilin, grown overnight at 37℃. For expression, 500 μL of this overnight culture was inoculated into terrific broth (TB) medium (50 mL, containing 100 μg/mL ampicilin) and incubated at 37℃ and 180 rpm in a flask until the optical density at 600 nm (OD600) reached 0.6∼0.8. Protein expression was induced by the adding 1 mM Isopropyl β-d-1-thiogalactopyranoside (IPTG) and incubating at 16℃ with shaking at 180 rpm overnight.

The culture was centrifuged at 4,000 rpm for 10 min at 4℃ to pellet the cells. The supernatant was discarded, and the pellet was resuspended in Tris-HCL buffer (100 mM, pH 7.5) supplemented with 1 mg/mL lysozyme (Takara Bio). The mixture was incubated at 30℃ for 30 min and then cooled to 0℃ on ice. Cells were disrupted by ultrasonication using Ultrasonic Homogenizer JY92-IIN (Scientz) at 100 W with a cycle of 3-second pulse and 5-second pause) for 20 min. The cell debris was removed by centrifugation at 16,000 rpm for 20 min at 4℃. The supernatant was collected for protein purification.

The soluble recombinant MDHs were purified using Ni-NTA Agarose (Qiagen). The supernatant was loaded onto the column and washed with a wash buffer (100 mM Tris-HCL buffer, pH 7.5, 300 mM NaCl, 25 mM imidazole, 5% glycerol). Proteins were eluted with an elution buffer (100 mM Tris-HCL buffer, pH 7.5, 300 mM NaCl, 333 mM imidazole, 5% glycerol). The eluent was desalted using BeyoDesalt™ G-25 Midi Desalting Column (Beyotime Biotechnology) and 100 mM Tris-HCL buffer. The concentration of the desalted protein solution was determined using a NanoDrop 2000 system (Thermo Fisher Scientific).

To test for MDH activity, a reaction mixture (50 μL) in 100 mM Tris-HCL buffer containing 1 mM oxaloacetic acid and 1 mM NADH was prepared in a UV-transparent 96-well with a clear flat bottom (Beyotime Biotechnology). The reaction was initiated by adding 50 μL of purified protein to the mixture. The reaction was carried out at room temperature, and NADH oxidation to NAD+ was monitored by measuring absorbance at 340 nm (ε = 6.22 mM^-1^ cm^-1^) every 30 seconds for 30 minutes using an Infinite® 200 PRO plate reader (Tecan). All reactions were performed in triplicates, and a mixture without purified protein served as the negative control.

For the biotransformation assay, the reaction mixture was scaled up to 250 μL and incubated at 25℃ overnight. All the reactions were performed in triplicates, and the mixture without purified protein served as the negative control. The biotransformation was quenched by adding equal volume of acetonitrile, followed by thorough vortexing and centrifugation at 12,000 rpm for 1 min at room temperature. The supernatant was filtered through a 0.22 μm ultrafitration membrane (Millipore), and malic acid was analyzed by HPLC-ESI-MS/MS on a 6475 Triple Quadrupole LC/MS System (Aglient). Chromatographic separation was achieved on an Acquity UPLC HSS T3 Column (1.8 µm, 2.1 × 150 mm; Waters) using a 1290 Infinity II ultra performance liquid chromatography (UPLC) system (Agilent). The separation was performed at 30℃ with an isocratic elution of 90% water (with 0.1% formic acid) and 10% methanol at a flow rate of 0.15 mL/min. The mass spectrometer was operated in negative ion mode with multiple reaction monitoring (MRM), using 133 *m/z* and 115.1 *m/z* as precursor and product ions, respectively. Fragmentation voltage was set to 60 V, collision energy to 5 V, gas temperature to 300°C, gas flow to 5 L/min, sheath gas pressure to 11 L/min, capillary voltage to 3500 V, and nozzle voltage to 1000 V. Data acquisition and analysis were performed using Agilent MassHunter Qualitative Analysis 10.0 (Agilent).

## Results

### Overview of the DNPD framework

We developed a convenient one-stop solution (Figure 1) for general-purpose protein design by integrating a state-of-the-art protein generative model, ProtGPT2 ^11^ with a custom-built protein discriminator. The ProtGPT2 model, a large-scale language model with 738 million parameters, serves as the foundation for generating new protein sequences. The protein discriminator, consisting of a convolutional neural network followed by a fully connected neural network (Figure 1A), is used to assess and prioritize the generated sequences. Protein sequence features are extracted using ProtT5-XL-UniRef50, a model-based embedding framework in ProtTrans collection, which comprises of 3 billion parameters ^39^. The generative model is finetuned with a positive set of protein sequences, while the discriminator is trained on a balanced dataset containing both positive and negative protein sequences. As proof of concepts, we applied this framework to the design of two distinct protein classes: antibacterial peptides (AMP) and malate dehydrogenases (MDH; EC1.1.1.37).

### Discriminators accurately classifies AMP/non-AMP and MDH/non-MDH sequences

For the AMP/non-AMP discriminator, the AMP and non-AMP sequences were obtained from SATPdb database ^33^ and UniProt ^34^, respectively. Short peptides from UniProt that are shorter than 50 residues and without certain antimicrobial-relevant keywords were selected as non-AMPs ^35^. Similarly, for the MDH/non-MDH discriminator, we acquired MDH sequences from Repecka et al. ^21^ and non-MDH sequences randomly sampled from UniProt ^34^. To ensure unbiased results, all training, validation, and test sets were carefully balanced by randomly sampling negative sets to mirror the sequence length distribution in positive sets. The discriminators achieved high accuracies and converged rapidly (<100 epochs), demonstrating the effectiveness of transfer learning-based embedding techniques. Specifically, the AMP/non-AMP discriminator achieved an accuracy of 0.81 on a balanced testing set of 576 samples (Figure 2A), while the MDH/non-MDH discriminator attained an accuracy of 0.99 on a larger balanced test set of 28,412 samples (Figure 2A). It is important to note that numerous AMP/non-AMP discriminators have been developed by other researchers ^55^. However, our framework is meant for de novo design for peptides/proteins in general, instead of building a discriminator for a specific task such as classifying AMP.

**Figure 2.**
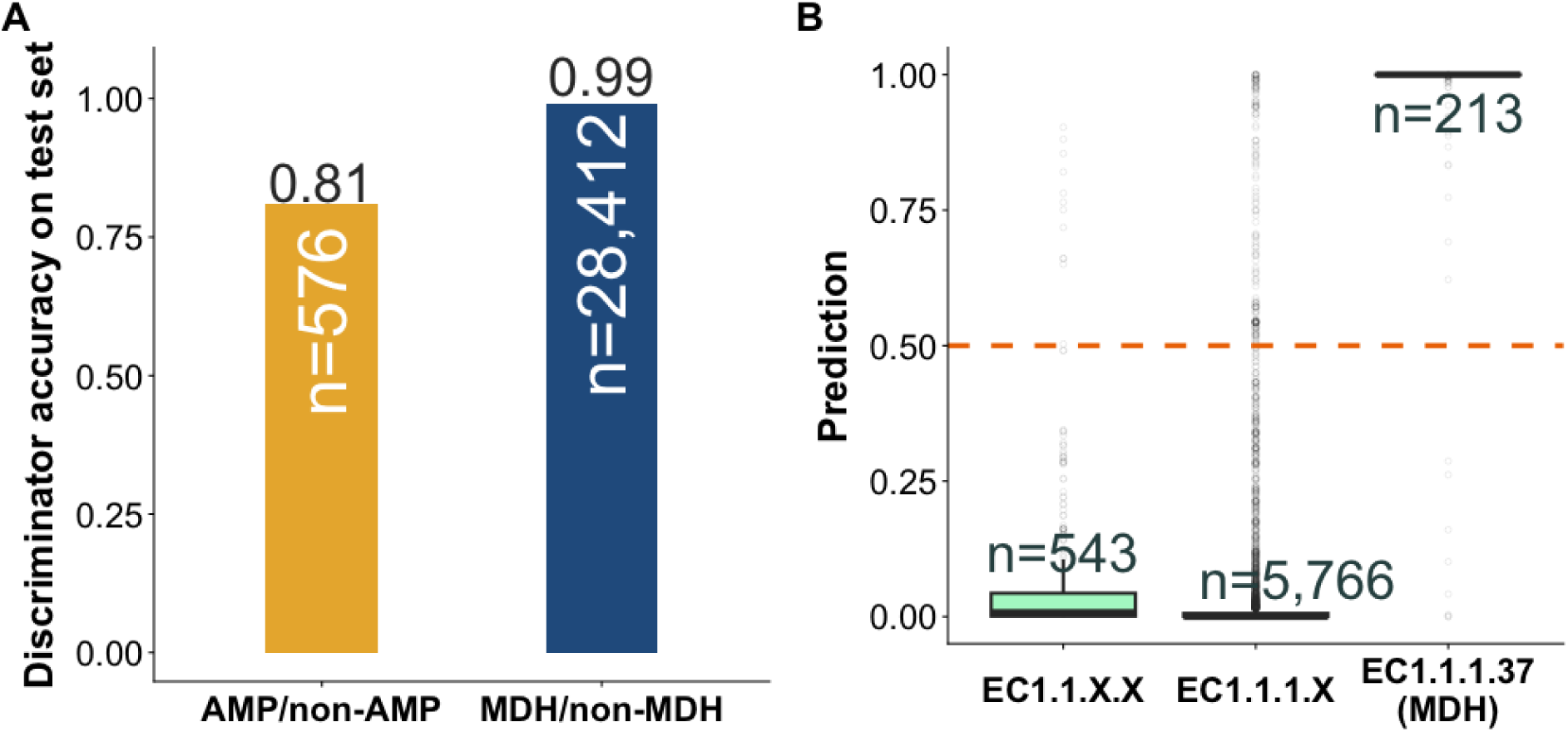
Accuracies (A) and resolutions (B) of the discriminators. In panel A, numbers denoted inside the bars are size of testing set (both balanced). Panel B shows MDH/non-MDH discriminator’s predictions on MDHs and functionally close enzymes: “EC1.1.1.X” are enzymes with the first three EC digits being 1.1.1. (excluding EC1.1.1.37); likewise, “EC1.1.X.X” are enzymes with the first two EC digits being 1.1. (excluding EC1.1.1.X). Numbers above the boxes are data size, and all data are not included in training set. AUCs for EC1.1.X.X vs. MDH and EC1.1.1.X VS. MDH are both 0.99.

To further evaluate the discriminators’ capability to differentiate between functionally close enzymes, we assembled additional datasets comprising EC1.1.1.X enzymes (n=5,766, excluding EC1.1.1.37) and EC1.1.X.X enzymes (n=543, excluding EC1.1.1.X), as well as an additional reserved set of 213 MDHs for further examination. All of these sequences were unseen during training. The MDH/non-MDH discriminator achieved a 0.99 ROC-AUC for both EC1.1.1.X vs. MDHs and EC1.1.X.X vs. MDHs (Figure 2B), highlighting its exceptional ability in differentiating between functionally similar enzymes. Overall, the results demonstrated the effectiveness of the discriminators in accurately distinguishing protein sequences of interest.

### Finetuning increases the likelihood of the generated sequences being AMP or MDH

Finetuning essentially involves adapting a pre-trained LM to a custom input dataset, enabling the updated LM’s to generate content more closely aligned with the input’s style ^56^. We finetuned ProtGPT2 with 2,879 known AMPs and subsequently generated 3,000 AMP candidates using the finetuned model. During generation, the “maximal length” was set to 20 and no starter sequence was provided, however, it was to be noted that the actual generated sequences may exceed the set “maximal length”. For comparison, we created an equal amount of a) random sequences, i.e., randomly constructed sequences following the length distribution as natural AMPs; b) “AA-imitating”, i.e., following natural AMPs’ amino acid and length distribution; c) ProtGPT2-random sequences, i.e., sequences generated by the non-finetuned ProtGPT2. As expected, the finetuned-generated sequences are more likely to be actual AMPs than “AA-mimicking” and “ProtGPT2-random” (prediction median 0.87 vs. 0.37 and 0.01, Wilcoxon signed rank test p-value<2.2e-16; Figure 3A). Surprisingly, we noticed that the “AA-mimicking” group also reached moderately high prediction (median=0.37; Figure 3A). This can be attributed to two reasons—a) there lacks gold standard negative set and our positive set is relatively small (excluding anti-yeast, anti-viral, etc.); and b) cationic charge and hydrophobicity are the dominant determinants of antimicrobial effects ^57–59^. Since these “AA-mimicking” peptides closely mimics the amino acid profile of natural AMPs, they inherit the cationic charge and hydrophobicity and are thus more likely to possess antimicrobial effects.

**Figure 3.**
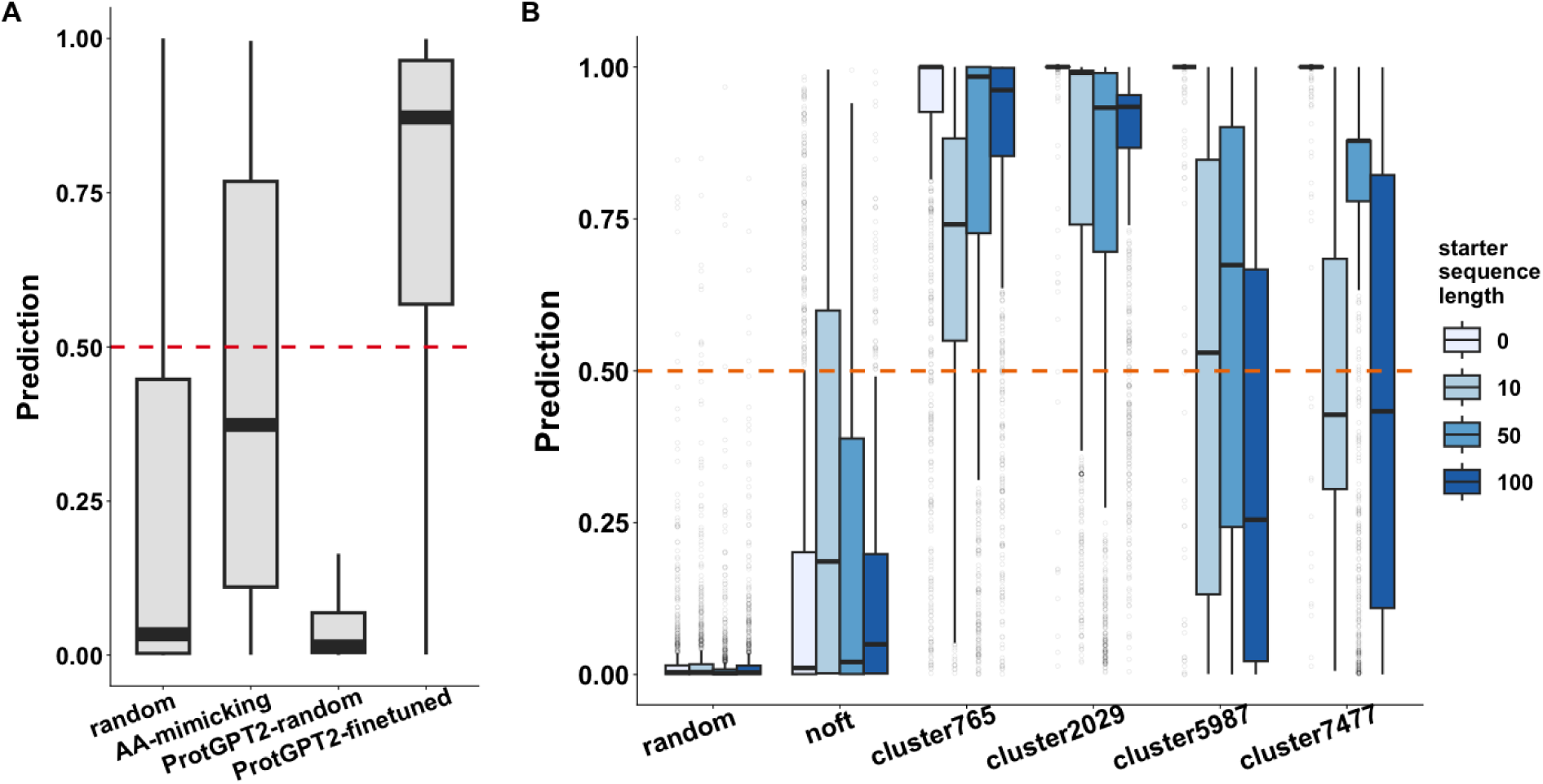
Effects of generator finetuning on AMP (A) and MDH (B) likelihood of the generated sequences. Panel A shows the discriminator’s predictions on four groups of peptide sequences. Group “random” means randomly constructed peptides; group “AA-mimicking” means peptides constructed following the amino acid and length distribution of the actual AMPs; group “ProtGPT2-random” means peptides generated by ProtGPT2 without any finetuning; group “ProtGPT2-finetuned” means peptides generated by ProtGPT2 that was finetuned with actual AMPs. In panel B, group “random” means random sequences following amino acid distributions MDHs; group “noft” in panel B means candidates generated by non-finetuned ProtGPT2. Other groups in panel B (*cluster765*, *cluster2029*, *cluster5987* and cluster7377) indicates sequences generated by ProtGPT2 finetuned by the respective sequence cluster (the numbers indicate corresponding cluster size).

To explore the impact of data size and sequence redundancy on finetuning results for MDHs, we clustered the 16,706 known MDHs and selected the four largest clusters (termed *cluster765*, *cluster2029*, *cluster5987*, and *cluster7477*) as finetuning sets, comprising 765, 2029, 5987, and 7477 sequences, respectively. As controls, we generated two sets of sequences: a) random sequences that followed the amino acid and length distribution of the known MDHs; and b) sequences generated by the non-finetuned model. Discriminator predictions revealed that sequences generated by the finetuned models were significantly more likely to be MDHs than random and non-finetuned model-generated sequences (prediction median 0.91 vs. 0.003 and 0.91 vs. 0.033, respectively; Wilcoxon signed rank test p-value<2.2e-16 for both comparison; Figure 3B). Notably, we found that even a relatively small number of sequences (∼700, in the case of cluster765) were sufficient to fine-tune ProtGPT2 for generating high-quality MDH sequences. Importantly, the discriminator’s predicted scores do not correlate with ProtGPT2’s perplexity during generation (Pearson correlation= -0.01). Considering the aforementioned limitations of perplexity, we further demonstrated the superiority of our discriminator in candidate prioritization.

Note that users can choose to provide a leading sequence to prompt ProtGPT2 during sequence generation, which we refer to as “starter sequence”. To investigate whether such a starter sequence would guide the generation towards our desired functions, we incorporated starter sequences of variable lengths. We supplied the first 0 (i.e., none), 10, 50, or 100 residues of a representative sequence (the representative sequence in the corresponding cluster; see Materials and Methods) as a starter sequence during the generation process. As a result, providing a starter sequence did not necessarily improve the quality of the generated sequences, as candidates generated without starter sequences generally had higher discriminator predicted scores (Figure 3B). Notably, in *cluster5987 and cluster7477, employing a starter sequence adversely impacted prediction accuracy. This suggests that, for a diverse training set, the inclusion of a starter sequence might impose unnecessary constraints on sequence generation*.

We observed that most of the generated sequences were non-redundant identified by cd-hit ^37^, except for those generated by the *cluster2029*-finetuned model (ratio of non-redundant sequences =0.31 vs. 0.67-0.95 in other generated groups), likely due to the low non-redundancy within the cluster itself (ratio of non-redundant sequences =0.08 vs. 0.28-0.33 in other clusters; Figure 4A). The effect of supplying a starter sequence on the redundancy of generated sequences was not conclusive (Figure S1).

**Figure 4.**
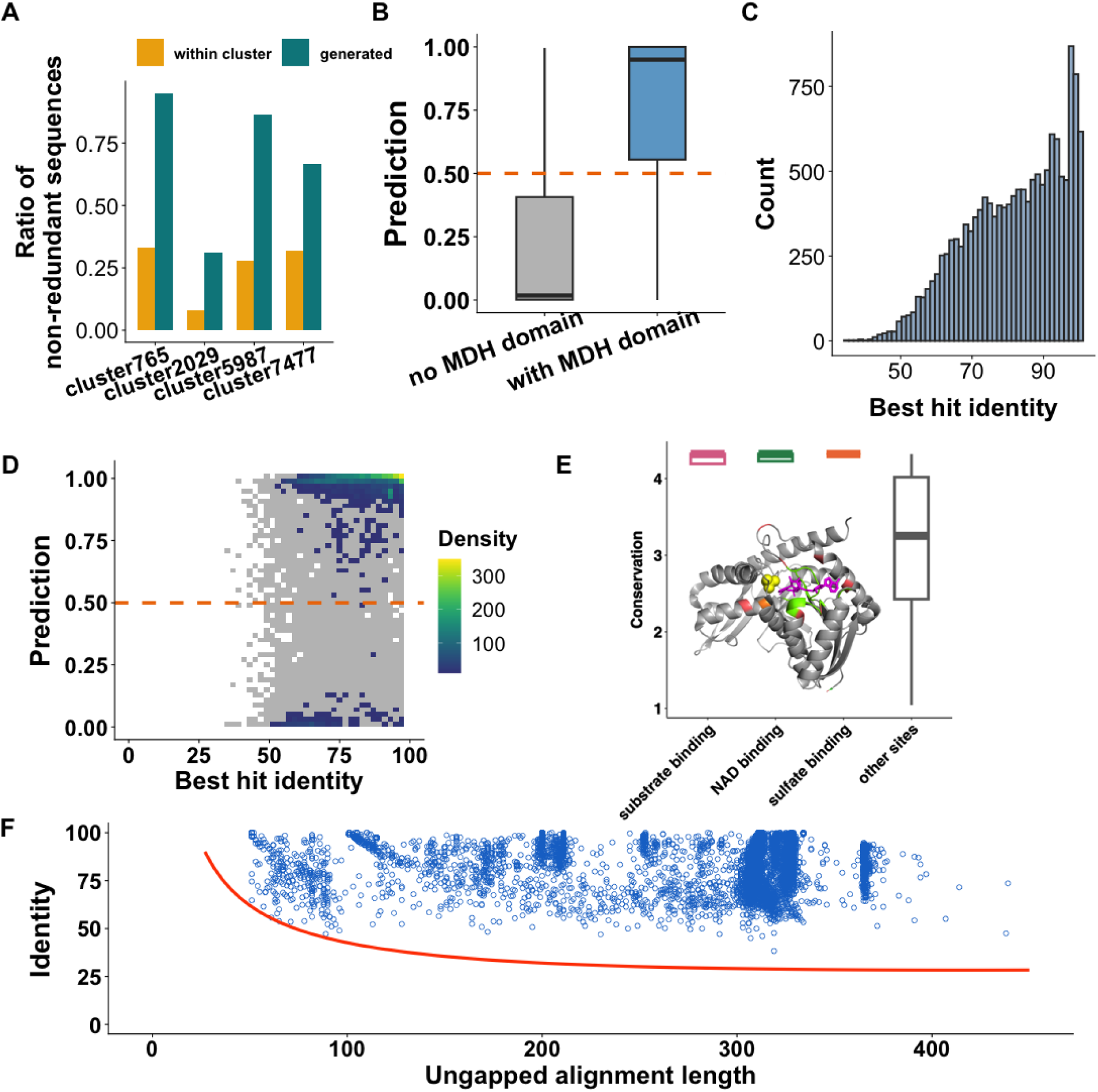
Computational analysis and evidence for the functionality of the prioritized MDH candidates. Panel **A** shows the ratios of non-redundancy within each cluster (*cluster765*, *cluster2029*, *cluster5987* and cluster7377) and among the generated sequences finetuned by the corresponding cluster. Panel **B** shows discriminator predictions on generated MDH candidates with and without MDH domains. Panel **C** shows the histogram of best hit identity of the generated MDH candidates. Panel **D** shows discriminator’s prediction vs. best hit identity (presented as data point density, grey grids contain <10 data points) for the finetuned-generated MDH candidates. Panel **E** shows the conservation along the MDH candidates with respect to functionally important sites. Position-specific conservation scores were derived from multiple sequence alignment (MSA) among 500 sequences (a known MDH sequence [UniProt ID: P11708] and 499 randomly sampled finetuned-generated MDH candidates with discriminator prediction>0.95). The determined protein structure (PDB ID: 4MDH) of P11708 was added to the MSA to map the protein positions to functionally important sites. The structure image depicts the structure of 4MDH with ligands sulfate (yellow spheres) and NAD (purple licorie) highlighted. Resides highlighted in pink, green, and orange indicate substrate binding, NAD binding, and sulfate binding sites, respectively. The boxplot summarizes the conservation scores at different sites. Panel **F** shows the best hit’s identities and ungapped alignment lengths of generated sequences (blue dots) with high predicted scores (>0.95). Red curve in panel E indicates HFSP curve: sequence pairs above and below the curve are predicted by HFSP as having the same and different enzyme functions, respectively.

### Prioritized generated AMP candidates are endowed with antibacterial activity

With the generated AMP candidates and their corresponding discriminator predictions available, we randomly selected 24 AMP candidates with predicted scores greater than 0.95; meanwhile, 10 random peptides were generated as control (Table S1). The amino acid and peptide length distributions of the random peptides are the same as the natural AMPs’. These peptides were tested against six different bacterial strains, including four Gram-positive strains (*S. aureus*, *K. rhizophila*, *B. pumilus*, and *B. subtilis*) and two Gram-negative strains (*E. coli* GDMCC 1.335 and DH5α). Out of the 24 generated AMP candidates, six completely inhibited the growth of at least one bacterial strain (Figure S2). Notably, five of these candidates displayed broad-spectrum activity against all six bacterial strains. In comparison, only one of the randomly selected peptides exhibited activity against some bacterial strains, highlighting the presence of many unknown AMPs in nature, as reported in previous studies ^60,61^. To accurately assess the potency of these candidates, five broad-spectrum AMPs were purified to 90% purity and their minimum inhibition concentrations (MICs) were determined by a growth-based assay. Four of these candidates displayed remarkably low MICs (with the lowest at 1.875 μM, Table 2), highlighting their potential for therapeutic and industrial applications.

**Table 2.**
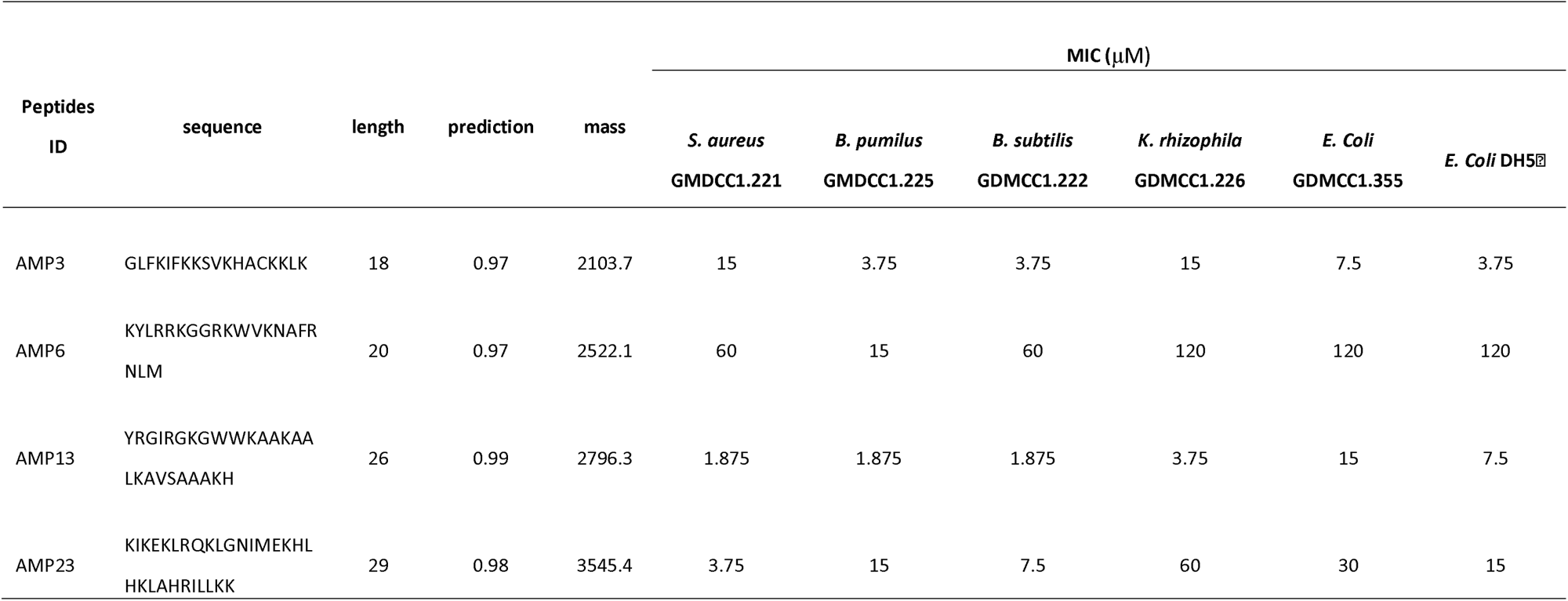
MIC of the identified broad-spectrum AMPs.

### Most finetuned-generated MDH candidates harbor MDH domains

A domain is a structural unit that carries a certain protein function. The function of MDHs is characterized by a “lactate/malate dehydrogenase” or a “Malate/L-lactate dehydrogenase” domain (here we denote as “MDH domain”) catalogued in Pfam database ^62^. Thus, bearing such signature domain serves as a good indicator of generation quality in addition to our discriminator. We found that not a single random or non-finetuned-generated sequence has the MDH domain (Figure S3), suggesting that this domain is extremely unlikely to occur by chance. In contrast, the vast majority (94.54%) of finetuned-generated sequences have MDH domains, increasing the confidence in their potential to possess the desired MDH function. Besides, the discriminator’s predictions on sequences with MDH domains are significantly higher than those without MDH domains (prediction median 0.95 vs. 0.02, Wilcoxon signed rank test p-value<2.2e-16; Figure 4B), assuring that the discriminator can well capture the sequence-function relationship.

### Homology inference suggests MDH function in most finetuned-generated MDH candidates

The MDH candidates generated through fine-tuning exhibit high sequence identity (median=83.65%) with their best hit known MDH (Figure 4C). Meanwhile, we believe that some high identity candidates may not actually be MDH (Figure 4D, bottom right corner) according to our discriminator. In contrast, some candidates with relatively low best hit identities (50-60%) exhibit high predicted scores, enabling researchers to select MDH candidates with a relatively high degree of novelty. Next, to further validate the utility of the discriminator in light of homology, we selected a set of generated candidates highly likely to be MDH, i.e., prediction>0.95, defined as *prioritized candidates*. We applied HFSP (homology-derived functional similarity of proteins) ^47^ to the *prioritized candidates*. HFSP assesses whether a pair of proteins share the same function based on ungapped alignment length and alignment identity. Although resolution of HFSP may not be able to reach the fourth level of EC number ^47^, it is still a useful baseline for homology reference. Notably, HFSP predicted that all prioritized generated candidates have the same function as their corresponding best hit sequence (Figure 4F), i.e., a known MDH. These results reinforced our confidence in the quality and functionality of the MDH sequences generated through fine-tuning, as supported by homology analysis.

### Prioritized MDH candidates are close to known MDHs in sequence space

To visualize spatial relationships between sequence-derived features of the known MDHs and the discriminator-prioritized MDH candidates, we first embedded the protein sequences into L×1024-dimensional vectors (where L is the length of a protein) using ProtTrans ProtT5-XL-UniRef50, we than used t-SNE algorithm ^48^ to transform the vectors into a two-dimensional space. Here, we randomly selected 100 known MDHs and 100 *prioritized candidates* to be visualized. To avoid bias caused by redundant sequences, we further limited the *prioritized candidates* to be non-redundant with each other determined by cd-hit ^37^ during the random selection. Functionally similar enzymes (EC1.1.1.X and EC1.1.X.X) were also added into the comparison as reference. The resulting visualization (Figure S4) showed that known MDHs and the *prioritized candidates* tend to cluster together, while functionally similar enzymes barely locate in the space between the known MDHs and the prioritized MDH candidates. This observation suggests that, in terms of sequence context, the *prioritized candidates* are in line with the known MDHs and are distinct from functionally similar enzymes.

### Prioritized MDH candidates are conserved at important functional sites

We randomly selected 499 *prioritized candidates* to perform multiple sequence alignment (MSA) with a known MDH with structural information (UniProt ID: P11708; PDB ID: 4MDH). From the MSA, we derived position-specific conservation scores and mapped them to 4MDH’s structural annotations. We found that the conservation scores at substrate binding sites, NAD binding sites, and sulfate binding sites were significantly higher than those at other sites (11 vs. 7, t-test p-value<0.00025 for all three comparisons; Figure 4E). These results suggest that our framework has successfully learned the sequence patterns required for MDH function and that the resulting MDH candidates are likely to perform this function.

### Prioritized MDH candidates are functional in vitro

We randomly selected 10 prioritized (predicted score > 0.9) and novel MDH (< 50% identity to any natural MDH) candidates generated from our framework for experimental validation. As positive controls, we randomly selected 10 natural MDHs from the training set (n=16,705) of ProteinGAN ^21^. Information of these sequences is summarized in Table S2. We expressed these proteins in *E. coli* BL21 (DE3) system (Figure S5). We observed that only 3/10 MDH candidates and 4/10 positive controls were soluble (Figure 5 A). Activity assays showed that all the soluble MDH candidates and the positive controls are functional in vitro (Figure 5 B-C), showcasing the utility of our framework in designing functional and novel enzymes.

**Figure 5.**
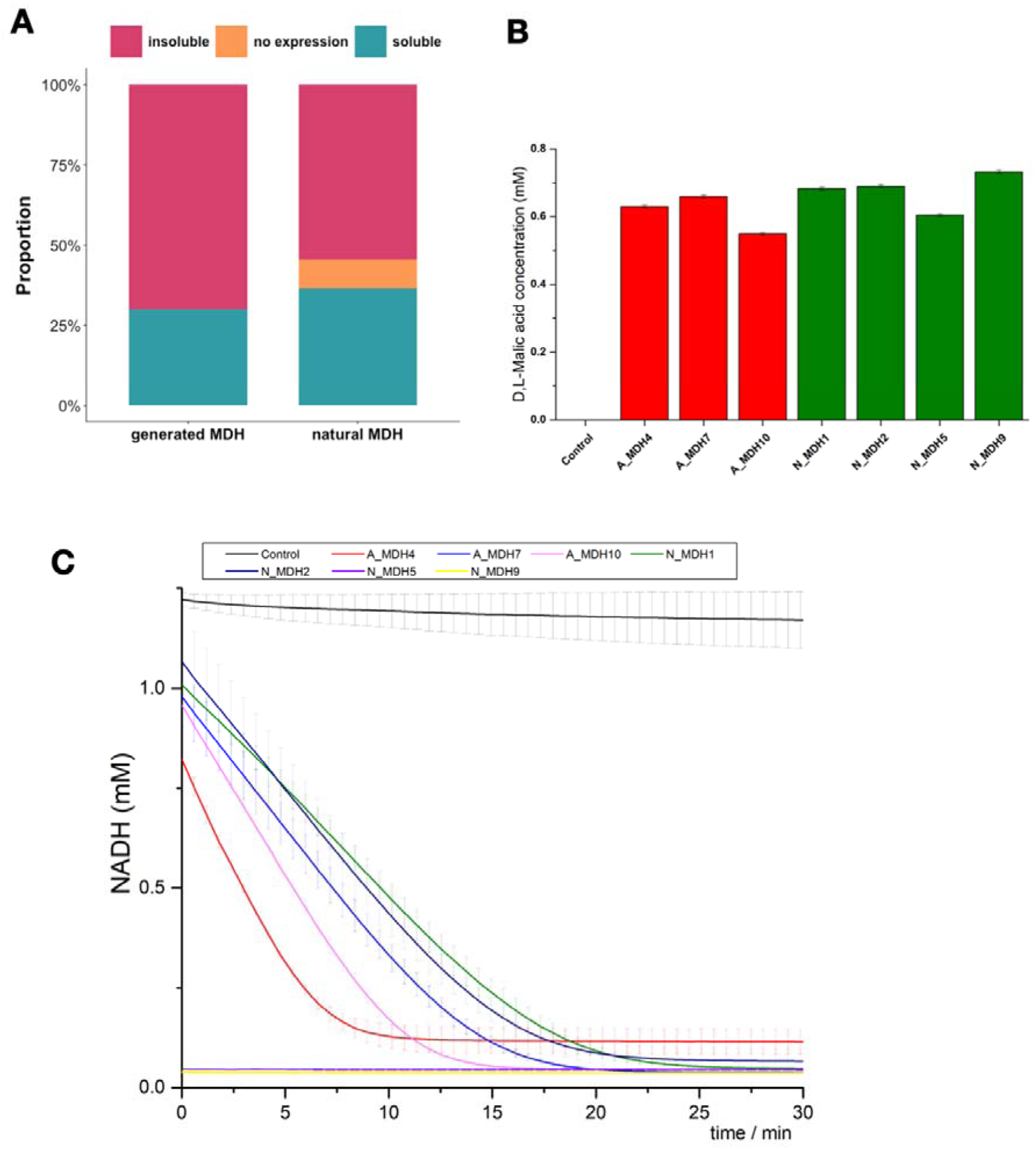
Experimental validation on the generated and prioritized MDH candidates. Ten prioritized (prediction > 0.9) and novel (<50% sequence identity to any natural MDHs) were randomly selected. Ten natural MDHs were also randomly selected as positive control. Samples without recombinant enzymes served as negative control. Results showed that 3/10 generated candidates and 4/10 positive controls were expressed and soluble (no significant difference; proportion test p-value=1), as displayed in panel A. Panel B shows the malic acid concentration transformed from oxaloacetic acid. Panel C shows the consumption of NADH over time. Prefix letter A and N indicates the artificially generated group and the natural group (positive control) group, respectively. For example, “A_MDH4” indicates the fourth generated MDH candidate and “N_MDH1” indicates the first positive control.

## Discussion

AMPs offer hopes in addressing the global antibiotic-resistance crisis ^63^ due to their advantages over small molecule antibiotics, e.g., modulation of immune response, broad-spectrum activity, multiple mechanisms of action, and diverse source organisms from all domains of life ^64^. Numerous attempts have been made to predict and *de novo* design novel AMPs ^65–67^. Recently, there are four deep generative language model-based methods for AMP design with experimental validation ^61,68–70^. The first three methods involved an RNN-based generator and a predictor, both trained on known AMPs. In addition to the predictor, a variety of physicochemical properties were also employed to facilitate candidate filtering. The fourth method ^70^ is more complex—an autoencoder generator was trained on all known short peptide sequences to better reflect peptide properties and enable high diversity. Meanwhile, multiple attribute classifiers (such as antimicrobial activity, toxicity, sequence similarity, and physicochemical properties) were trained on the latent space of the autoencoder, and the generation process was guided by these attribute classifiers altogether for desired peptide properties, and the attribute classifiers and a series of external predictors were used for candidate screening. All of these language model-based methods, as well as GAN-based methods ^71^, have successfully identified experimentally valid novel AMPs, some of which are broad-spectrum. However, performance comparison across these methods is difficult due to variations in data size, time consumption for training, threshold of “inhibitory effect”, definition of “broad-spectrum”, bacterial strains tested, peptide purity, and sequence novelty. From a usage convenience standpoint, our framework has two major advantages over other language model-based methods. First, it does not need to train a generator from scratch, as the existing generator (ProtGPT2) assumably performs better due to the tremendously larger training protein dataset. Second, the candidate prioritization is straightforward as it relies on only one discriminator, instead of multiple classifiers or additional calculation of physicochemical properties.

Enzyme engineering has become an essential aspect of metabolic engineering and synthetic biology ^72,73^ . When incorporating a heterogenous enzyme into an engineering chassis, it is often necessary to adjust the enzyme properties to better fit the working condition. Generating artificial protein sequences offers more options for selecting the ideal enzyme version. For example, a variety of tools can be applied to screen for more suitable solubility ^74,75^, catalytic capacity ^76,77^, and protein-protein interaction ^78,79^. Many attempts have been made for enzyme DNPD ^80,81^. One of the most successful non-LLM frameworks is ProteinGAN ^21^, a GAN-based generator. As a showcase, ProteinGAN was trained on 16,706 MDHs and produced 55 candidates for testing, of which 13 candidates experimentally displayed MDH catalytic activity. Notably, GAN is suitable for continuous data (such as images), but not for discrete data (such as text), because GAN relies on gradient-based optimization and there is no smooth gradient between distinct tokens ^82^. When applied in text (such as protein) generation tasks, GAN also suffers from training instability ^83^, ignorance on sequential dependencies ^84^, mode collapse ^85^, and result evaluation ^86^.

Compared to ProteinGAN, our framework has several advantages. First, our discriminator was trained on much less protein sequences than ProteinGAN (n=2,029 vs. n=16,706) without compromising the accuracy (0.99 accuracy on a much larger testing set: n=28,412). Second, it can distinguish MDHs from functionally similar enzymes at a high resolution, surpassing the limitations of traditional homology-based methods ^47^. The candidate screening process is therefore remarkably simplified and improved, eliminating the need for additional predictors and replacing sequence identity-based criteria with a straightforward sequence-function scoring scheme. With that said, it allows us to select both conservative (high sequence identity) and novel (low sequence identity) candidates while maintaining high functional fidelity (high discriminator scoring). Last but not least, our framework is time-efficient. The time consumption for the MDH task (in the case of the *cluster2029* for example) is dissected as follows: ∼10 min generator finetuning (n=2,029), ∼1 h embedding the training/validation sets (n=4,500 + 500), 2 min discriminator training/validation (n=4,500 + 500), ∼4 h generation (1,000 sequences), ∼20 min embedding the generated sequences (n=1,000), and ∼1 min prediction. The MDH task can be completed in 6 hours (without parallel computing), which is a small fraction of the time it took for ProteinGAN (9 days). Designing novel MDHs was also attempted in Madani et al.’s work on ProGen ^12^. In their work, the aforementioned per-token log-likelihood scoring, as well as ProteinGAN’s discriminator were used for the prioritization on the generated MDH candidates, achieving 0.94 and 0.87 ROC-AUC, respectively, on 56 samples obtained from the ProteinGAN paper ^21^. In comparison, our discriminator achieved higher ROC-AUC (0.99) on a much larger test set (n=28,412), demonstrating the accuracy of our discriminator.

A major limitation of our framework is that finetuning ProtGPT2 requires a large GPU memory: among the NVIDIA GPU hardware we have tried (A100, A10, V100, T4, P4, P100), only NVIDIA A100 (80G) was able to handle the workload. A minor disadvantage is that, unlike GAN requires only positive set, a negative set is also needed to train our discriminator. However, obtaining a negative set from protein sequence databases such as UniProt ^34^ is straightforward.

In summary, we developed a framework that can be quickly tailored to various peptide or protein tasks without the need to re-learn the protein grammar. Compared to other de novo peptide/protein design methods ^21,70,87^, our framework is time- and data-efficient, and is more convenient to use without the need to incorporating multiple filters or classifiers for screening purposes. Therefore, this framework may have the potential to significantly accelerate AI-based DNPD.

## Data and code availability

Codes for the framework and result analysis, as well as the data to replicate this work, were deposited in GitHub repository (https://github.com/zishuozeng/GPT_protein_design).

## Abbreviations used

GPT: generative pre-trained transformers
NLP: natural language processing
GAN: generative adversarial network
LM: language model
RNN: recurrent neural networks
AMP: antimicrobial peptide
MDH: malate dehydrogenase
EC: enzyme commission
MIC: minimum inhibitory concentration

## Acknowledgements

We would like to thank Yu Han (SIAT) for the constructive discussion and all researchers who made the data and tools relevant to this study available.

## Author contributions

Z.Z. and X.L. conceived the ideas; Z.Z. performed the modeling, data analysis, and result visualization; R.X. performed the AMP and MDH experimental validation and wrote the methods for it; Z.Z. wrote all other parts of the manuscript; J.G. performed preliminary finetuning experiments; X.L. revised the manuscript.

