## Supplementary Figure for "Binary Discriminator Facilitates GPT-based Protein Design"

**Supplementary Figures**

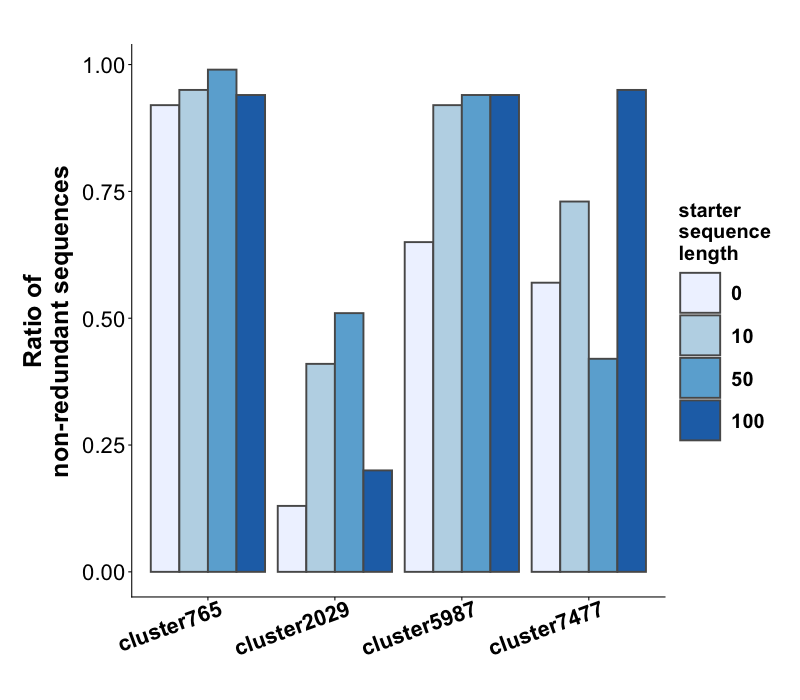

**Figure S1. Ratio of non-redundant sequences of the generated MDH candidates grouped by finetuned cluster and starter sequence length.**

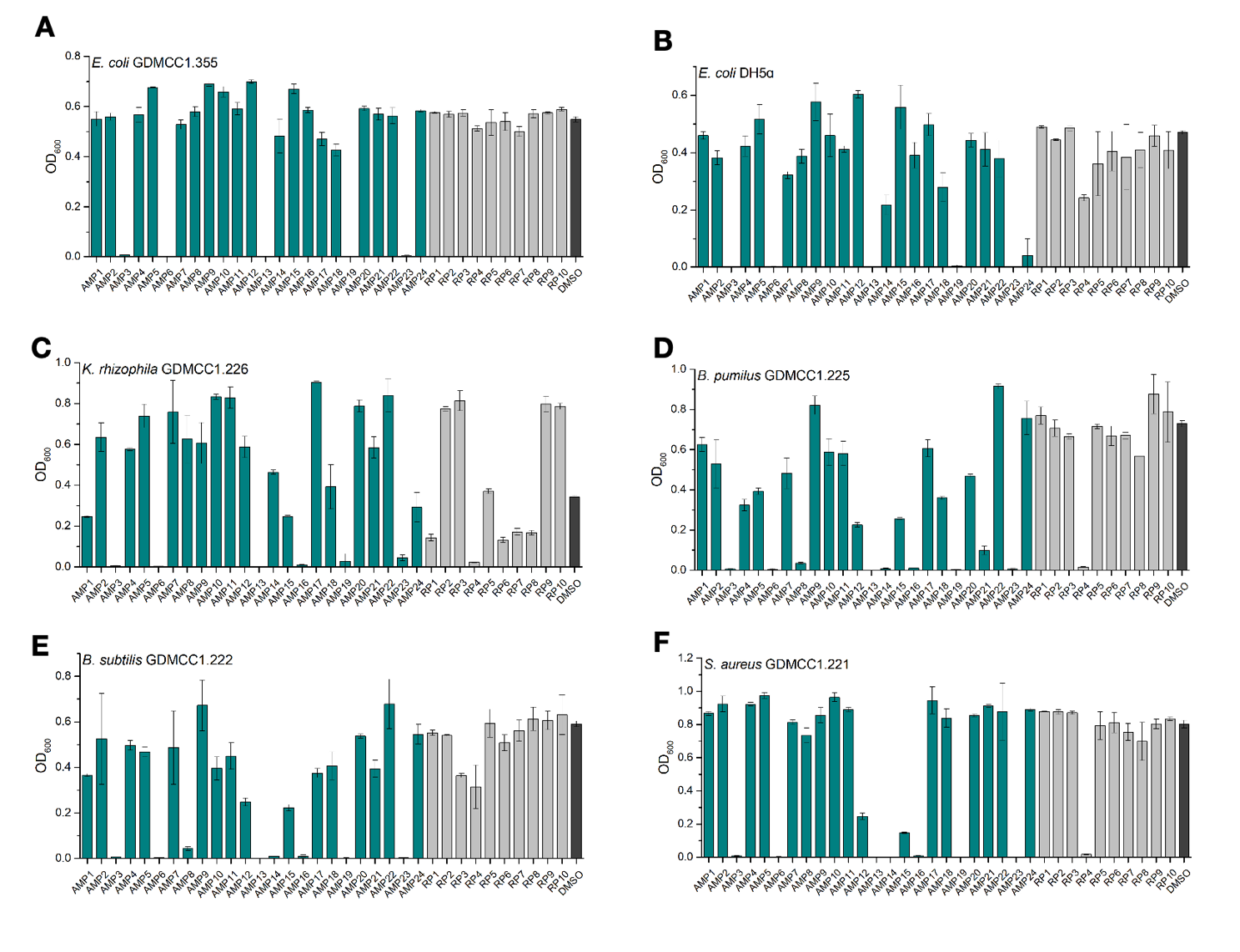

**Figure S2. Antibacterial effect of crude AMP candidates and random peptides.** “AMP1”-“AMP24” are AMP candidates, “RP1”-“RP10” are random peptides, “DMSO” is negative control. Panel A-F shows *E. coli* GDMCC1.355, *E.coli* DH4$\alpha$, *K. rhizophila* GDMCC1.226, *B. pumilus* GDMCC1.225, *B. subtilis* GDMCC1.222, and *S. aureus* GDMCC1.221, respectively. Lower OD_600_ value (Y-axis) indicates stronger antibacterial effect. Error bars are denoted for each group.

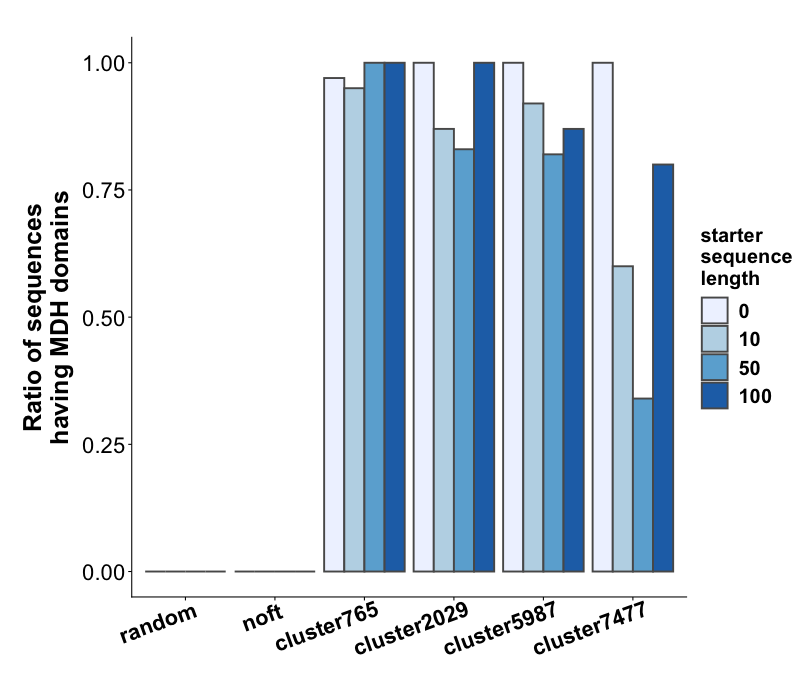

**Figure S3. Ratio of generated MDH candidates having MDH domain grouped by finetuned cluster and starter sequence length.**

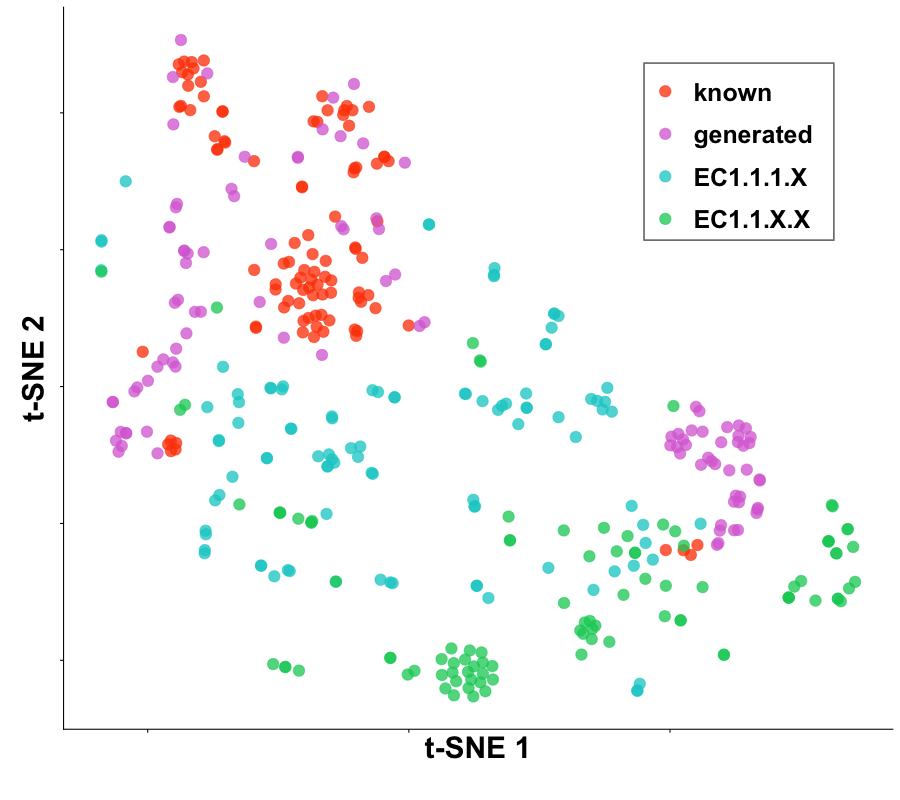

**Figure S4. Two-dimension t-SNE plot on sequences embedded as vectors.** The known MDH sequences (n=100) were sampled from *cluster2029*. The generated sequences (n=100) were sampled from MDH candidates generated by ProtGPT2 finetuned by *cluster2029* that are prioritized (non-redundant, discriminator prediction>0.95). Enzymes with EC number EC1.1.1.X (excluding EC1.1.1.37, i.e., MDH, n=100) and EC1.1.X.X (excluding EC1.1.1.X, n=100) were added to the plot to illustrate the relative spatial proximity between MDHs and functionally close enzymes. The t-SNE plot was generated based on protein numerical vector representation embedded by the ProtT5-XL-UniRef50 model.

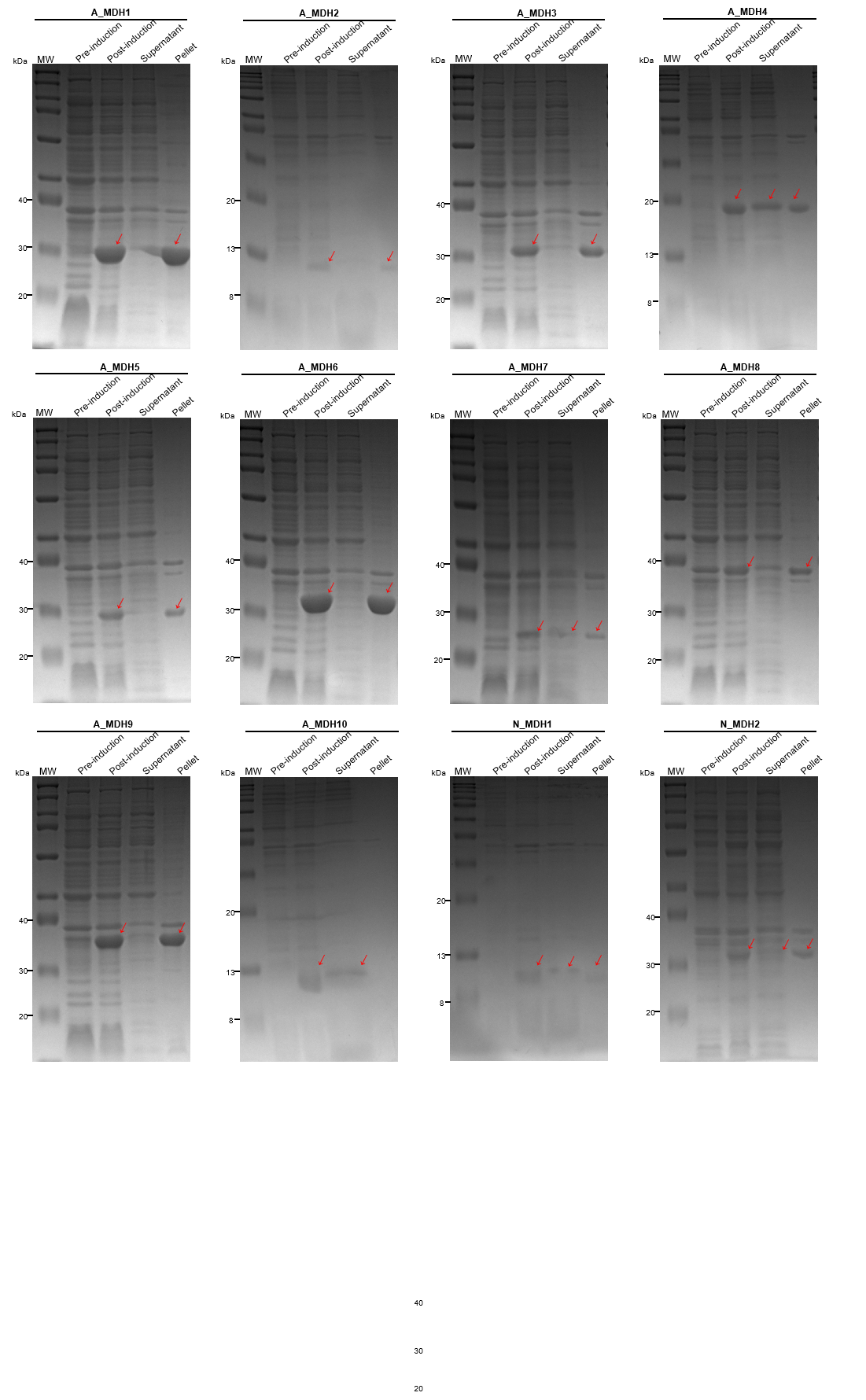

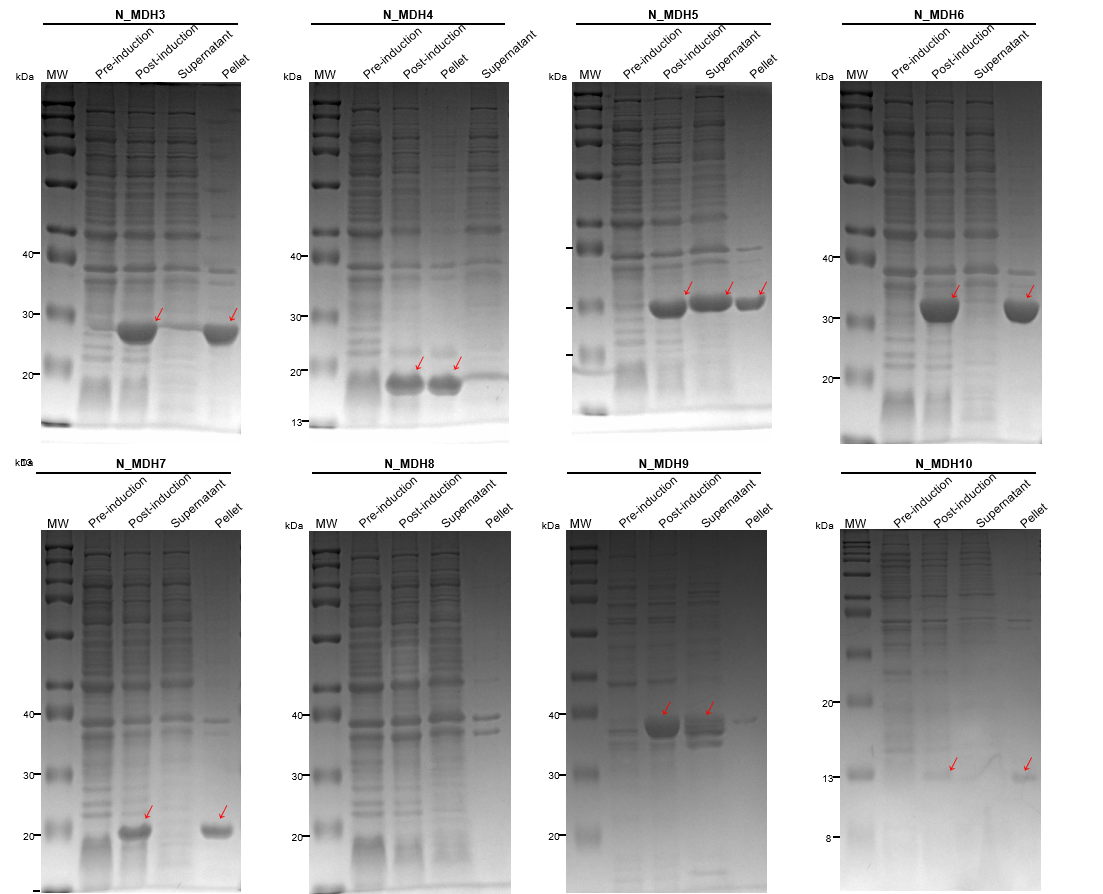

**Figure S5. Analysis of generated and natural MDHs overexpression in E. coli BL21(DE3) by SDS-PAGE**. The supernatant and pellet fraction of the lysate were also analyzed to verify the solubility of the proteins. Red arrow indicates the position where MDH is expected to be observed. Absence of red arrows indicates no expression. Prefix letter A and N indicates the artificially generated group and the natural group (positive control) group, respectively. For example, “A_MDH4” indicates the fourth generated MDH candidate and “N_MDH1” indicates the first positive control.
